# Hookworm genomic diversity and population structure from accessible sample types: A validated approach to generate genome-wide polymorphism datasets from individual third-stage larvae

**DOI:** 10.64898/2026.01.06.697970

**Authors:** Kaylee S. Herzog, Savanna Randi, Dickson Osabutey, Christina Paraggio, Richard Bungiro, Lisa Harrison, Irene Owusu Donkor, Francis Appiah-Tsum, Amanda Lamptey, Isaac Quaye, Shaelynn Vaughan, Michael D. Wilson, Anita Ghansah, Michael Cappello, Joseph R. Fauver

## Abstract

Hookworm infection is a neglected tropical disease affecting hundreds of thousands of people annually in the tropics and sub-tropics. Population genomic approaches have the potential to improve our understanding of hookworm infection dynamics and control efficacy. Here, we validate an approach to generate genome-wide polymorphism datasets from accessible sample types in the zoonotic hookworm (*Ancylostoma ceylanicum*) then apply the validated approach in the human hookworm (*Necator americanus*) to compare laboratory- and field-derived samples. We first present an optimized method for purifying nucleic acid from individual third-stage hookworm larvae (L3s). We then measure the accuracy of variant call datasets generated through whole genome amplification (WGA) and next-generation sequencing (NGS). We demonstrate that WGA via multiple displacement amplification (MDA) introduces predictable biases that are exacerbated by low inputs and poor sample preservation but show that with sufficient input mass (≥0.1ng) we are still able to produce highly accurate variant call datasets from nucleic acid concentrations that reflect those of individual L3s. Using our validated approach, we infer laboratory- and field-collected samples of *N. americanus* as distinct populations, with higher levels of heterozygosity and nucleotide diversity identified in field-collected samples, suggesting signatures of inbreeding and/or drift are detectable in laboratory specimens within several years of initiation of infection. We also show that, despite expected reductions in heterozygosity, laboratory samples still possess numerous heterozygous sites, and we demonstrate that a reference genome generated from an adult worm from an early laboratory passage performs well for variant calling in both laboratory- and field-derived samples. Moving forward, our optimized method for nucleic acid purification can be broadly applied to generate input for any amplification-based approach where sequencing individual hookworm L3s, rather than a pool of specimens, is preferred. Our validated population genomics workflow can be used to characterize structure and connectivity of hookworm populations in endemic communities, with the goal of leveraging these insights to improve our approach to hookworm treatment and control.

**AUTHOR SUMMARY:** Infection with the human hookworm, *Necator americanus*, is a significant cause of morbidity in the global south. Population genomic techniques have the potential to improve our understanding of hookworm infection and control. However, third-stage larvae, or L3, which are the hookworm life-cycle stage we routinely have access to when screening infected people, are less than a millimeter long, and this small size makes generating genomic datasets from individual worms difficult. Here, we introduce an optimized approach for purifying DNA from individual L3s and validate its use with whole genome amplification (WGA) for population genomics. We found that WGA from L3s produces sequencing datasets that are biased in their breadth and depth of coverage across the hookworm genome. But by ensuring a minimum threshold for the mass of DNA input to WGA and setting strict criteria for variant filtration, these biases can be overcome to produce highly accurate variant calls genome-wide. We then use this approach to demonstrate reduced genetic diversity in a recently established laboratory hookworm strain as compared to field-collected samples. Our study outlines a specimen-through-analysis workflow that can be used with accessible sample types to measure population structure and diversity of hookworms in endemic communities.

## INTRODUCTION

Hookworms are soil-transmitted helminths that blood feed as adults in the small intestine of their host, and billions of people each year are at risk for infection [1–3]. Though zoonotic species exist, most infections in humans are caused by two species: *Ancylostoma duodenale* and *Necator americanus*, with the latter being the most common [4–6]. Hookworm infection is considered a neglected tropical disease and causes significant morbidity, particularly in tropical and sub-tropical regions [7]. In these regions, hookworm infection is primarily controlled through preventative chemotherapy, with anthelminthic drugs distributed *en masse* to entire communities regardless of individual infection status [8]. Despite the prevalence of hookworm infection and the obvious harm it causes in vulnerable groups, little is known about how hookworm populations are assembled within endemic communities. Establishing baselines for hookworm population structure, diversity, and connectivity would allow for more robust evaluation of the impact of prophylaxis and treatment campaigns [9]. Population genomic approaches therefore have tremendous potential to improve not only our understanding of hookworm biology and infection dynamics, but also the efficacy of ongoing control efforts.

To date, the field of human hookworm population genetics has been largely based on small, targeted datasets in which one or two loci are sequenced, with a focus on *cytochrome oxidase subunit 1* (*COX1*) [10–13]. Only few studies have sampled a broader representation of the hookworm genome, including complete mitochondrial [14] and microsatellite [15] datasets. One of the major challenges in shifting hookworm population genetics into the next-generation sequencing (NGS) era is that the most accessible sample types are often small in size, making many NGS workflows untenable. For example, hookworm eggs and early-stage larvae can be obtained from both infected humans and laboratory animals, but at only tens of micrometers in size, sufficient nucleic acid input for NGS library preparation cannot be obtained without pooling individuals. Conversely, adult hookworms are large enough to generate genome-wide NGS libraries using nucleic acid from single worms [16], but obtaining adults from laboratory animals is a time-and labor-intensive process, and one that cannot be translated to infected humans. Third-stage larvae (L3) are therefore the most attractive stage in the hookworm life-cycle to target for population genomics. L3s are readily obtained from laboratory animals and from human subjects using non-invasive sample techniques (i.e., hatching and rearing hookworm larvae from fecal collections) [17]. At nearly a millimeter in total length, it is possible to separate individual L3s prior to nucleic acid extraction using a microscope, yet purifying a sufficient mass of genomic DNA (gDNA) for library preparation from one L3 remains a challenge.

Researchers conducting population genomic studies in other helminth systems have overcome the obstacle of low input sample types by using whole genome amplification (WGA). This approach, which commonly relies on multiple displacement amplification (MDA) to replicate template gDNA, has been leveraged for a variety of applications, including in the study of human disease [18]. In helminth population genomics, WGA has been applied to great effect; for example, in the blood fluke *Schistosoma mansoni,* using individual miracidia to investigate population connectivity and praziquantel resistance [19]; in the threadworm *Strongyloides stercoralis*, using individual L3s to elucidate population structure and transmission dynamics [20]; and in the filarial worm *Wuchereria bancrofti,* using individual microfilariae to assess whether post-treatment infections were due to reinfection or treatment failure [21]. In each of these examples, nucleic acid from a small yet accessible larvae stage was amplified using WGA to generate genome-wide polymorphism datasets and answer population-level questions. Given its successful application in other helminth systems, we sought to validate a similar approach for population genomics in human hookworms.

This study has two aims: (1) validate the use of WGA to generate genome-wide polymorphism datasets from individual hookworm L3s using an established laboratory strain of the zoonotic hookworm (*Ancylostoma ceylanicum*); and (2) apply the approach in the human hookworm (*N. americanus*) to compare laboratory- and field-derived samples. First, we optimized a technique for purifying nucleic acid from individual L3s. Next, we generated validation datasets by diluting nucleic acid from a single adult male of *A. ceylanicum* to represent the range of concentrations observed from L3 extractions. Then, we subjected these dilutions to WGA, library preparation, and NGS. We used genotype concordance metrics and principal components analysis (PCA) to compare single nucleotide polymorphism (SNP) datasets for each dilution against datasets generated from the same undiluted unamplified stock nucleic acid. Finally, we applied our validated approach to compare L3s of *N. americanus* from a more recently established laboratory strain to their contemporaneous counterparts from the same endemic region in Ghana originally used as the source population to initiate infection in laboratory animals.

## RESULTS

### Approach validation–Ancylostoma ceylanicum

#### Optimization of genomic DNA extraction from individual third-stage larvae

Cycle threshold values for nucleic acid extractions from individual L3s of *A. ceylanicum* using different lysis buffers are summarized in Figure 1. Of the three lysis buffers tested, the Zymo Quick-DNA^TM^ HMW MagBead Kit lysis buffer produced the highest-concentration extractions and the most consistent concentrations across samples (~0.01 ng/µL, or ~0.1 ng total per L3). This approach was therefore selected as the standard method for gDNA extraction from individual L3s throughout the study.

**Figure 1.**
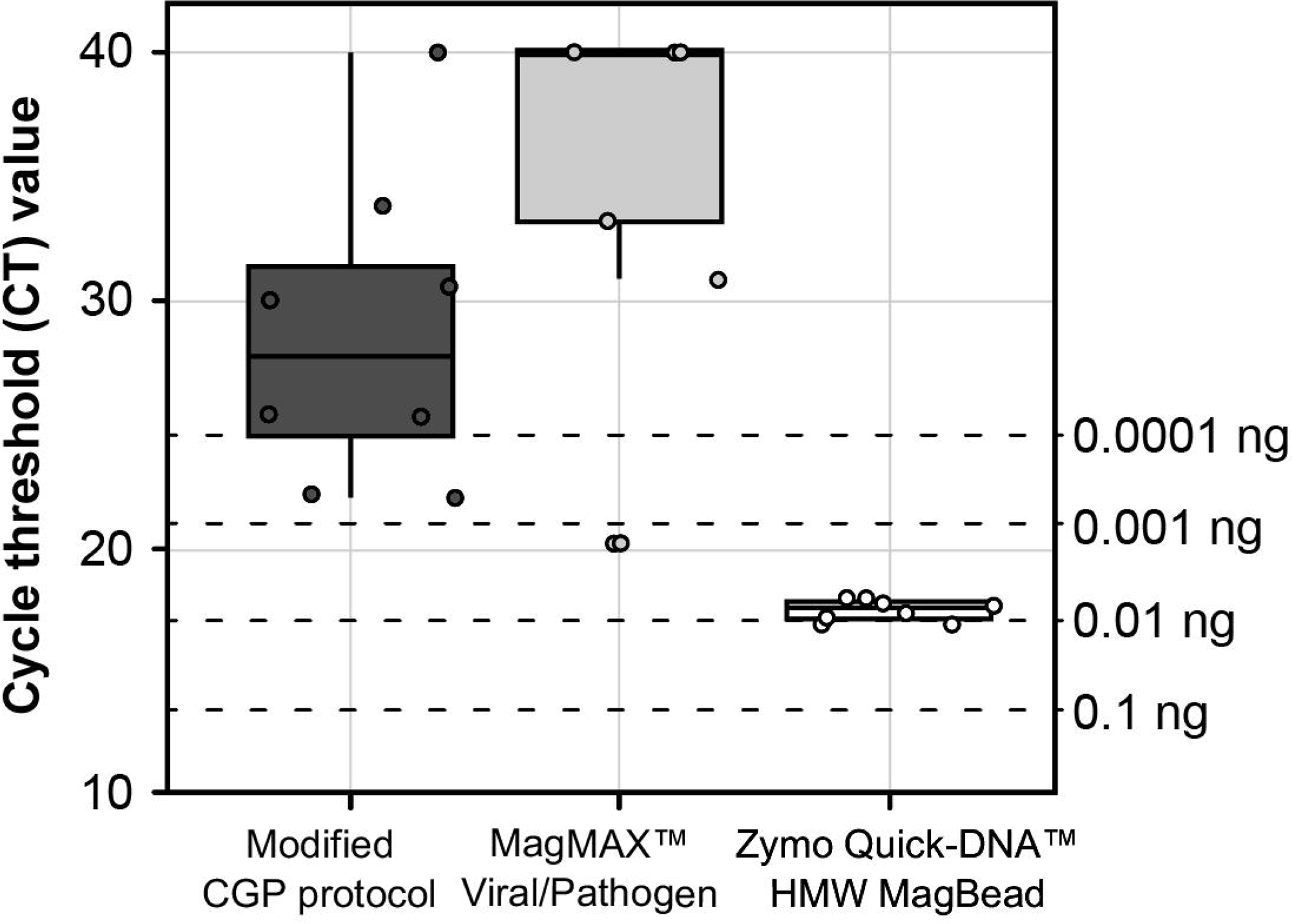
Extraction methods tested for individual third stage (L3) larvae of *Ancylostoma ceylanicum.* Ct values were generated using 1 µL of extracted genomic DNA as input to assay with the forward and reverse primers for ITSI of Llewellyn et al. (2016). Concentrations at right are obtained from standard curve generated from serially diluted genomic DNA from a single adult male worm.

#### Whole genome amplification, next-generation sequencing, and reference mapping

For the unamplified stock gDNA from the single adult male of *A. ceylanicum* (1ng/µL) and its amplified ten-fold serial dilutions (0.1ng/µL and 0.01ng/µL), inputs to WGA and library preparation and the number of reads generated and mapped to the reference genome are summarized in Table 1. Sample concentration post-WGA correlated positively with the total mass of nucleic acid input to amplification. For each dataset, ~97% of reads were retained after filtering by read length and quality. However, the proportion of reads that mapped to the reference genome in primary alignment and in proper pair varied linearly across datasets, with the highest rate observed in the unamplified stock gDNA (~95%) and the lowest rate observed in the lowest-concentration amplified dilution (~85%; 0.01ng input to WGA; see Table 1). Depth of coverage (DOC) plots for all three datasets are presented in Figure 2. A greater number of reads was needed for amplified dilution datasets to cover the same proportion of sites in the reference genome at >10× DOC as in the unamplified stock gDNA, and this disparity increased with decreasing input to WGA (Fig. 2A). When considering individual sites in at which reads mapped, DOC was more even for the unamplified stock gDNA as compared to its amplified dilutions. For the unamplified stock gDNA dataset, most sites (>80%) were covered by at 50× or greater, few sites (<5%) were covered at 100× or greater, and almost no sites were covered by at greater than 200× (Fig. 2B). For the higher-concentration amplified dilution dataset (0.1ng input to WGA) while most sites (>80%) were similarly covered at 50× or greater, >40% of sites were covered at 100× or greater, and nearly 10% of sites were covered at greater than 200×. For the lower-concentration amplified dilution dataset (0.01ng input to WGA), only ~50% of sites were covered at 50× or greater, and yet nearly 20% and 5% of sites were covered at 100× and 200× or greater, respectively (Fig. 2B).

**Figure 2.**
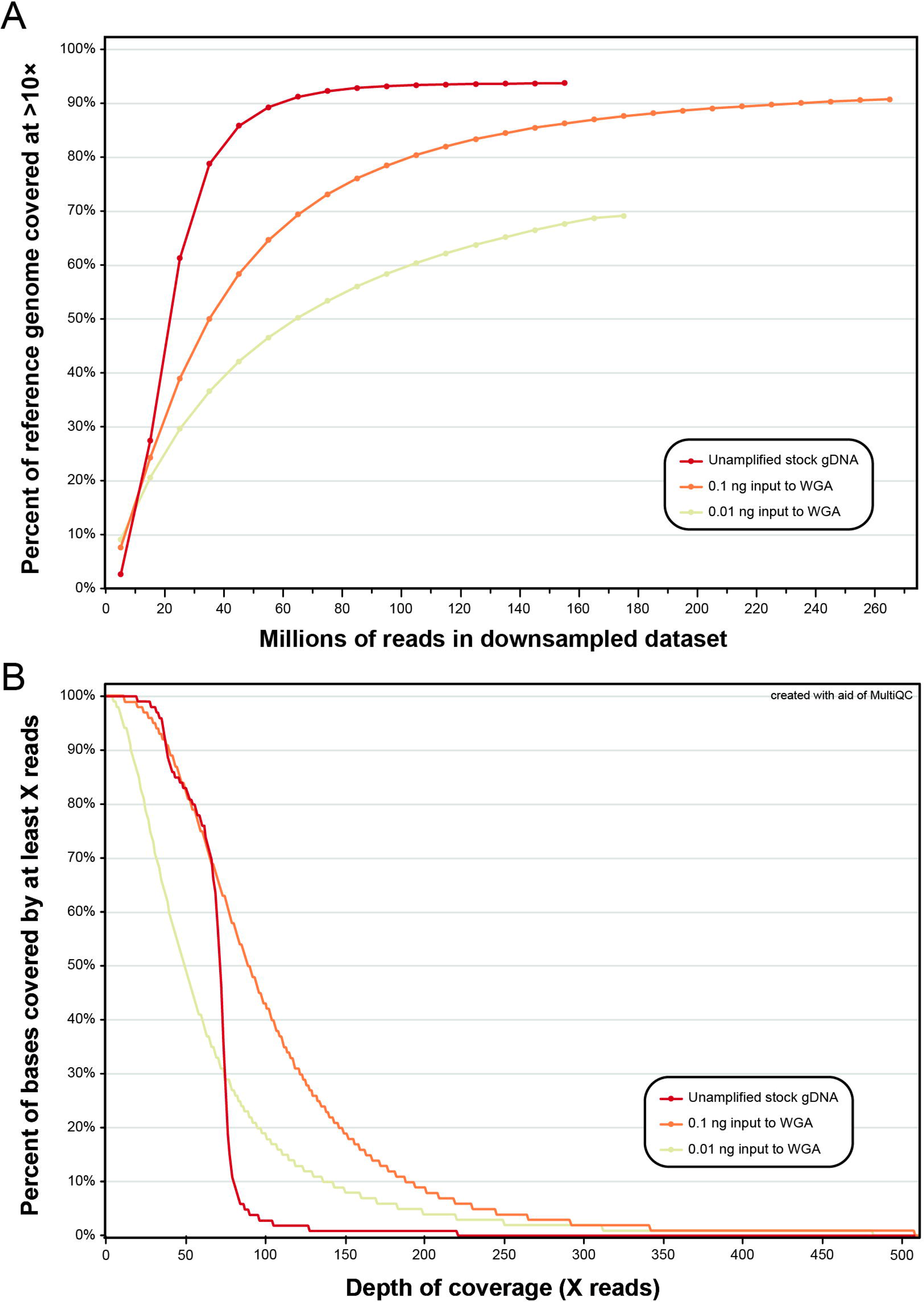
Depth of coverage plots for unamplified stock gDNA from a single adult male of *Ancylostoma ceylanicum* and its whole-genome-amplified (WGA) ten-fold serial dilutions. **(A)** Percent of sites in reference genome of Schwarz et al. (2015) covered at >10x depth of coverage versus proportion of reads in downsampled datasets. **(B)** Portion of sites in binary alignment map (BAM) files with reads mapped at a specified depth of coverage.

**Table 1.** Whole genome amplification (WGA), library preparation, next-generation sequencing and reference mapping statistics for stock genomic DNA from a single adult male of *Ancylostoma ceylanicum* and its serial dilutions. Reads were aligned to reference assembly of Schwarz et al. (2015).

|  | Unamplified<br>stock gDNA | 10<sup>-1</sup> dilution | 10<sup>-2</sup> dilution |
| --- | --- | --- | --- |
| **Concentration** | 1 ng/μL | 0.1 ng/μL | 0.01 ng/μL |
| **Input to qPCR** | 1 ng | 0.1 ng | 0.01 ng |
| **Average C<sub>t</sub> value** | 10.24 | 13.63 | 17.26 |
| **Input to WGA** | — | 0.1 ng | 0.01 ng |
| **Concentration post-WGA** | — | 54 ng/μL | 19.3 ng/μL |
| **Total mass post-WGA** | — | 1,080 ng | 386 ng |
| **Input to library preparation** | 3.8 ng | 250 ng | 146.4 ng |
| **No. cycles indexing PCR** | 10 | 5 | 5 |
| **No. reads generated** | 166,577,248 | 264,869,508 | 170,988,274 |
| **No. (proportion) of reads retained post-QC with fastp** | 162,538,778<br>(97.58%) | 257,414,872<br>(97.19%) | 165,682,332<br>(96.90%) |
| **No. (proportion) of reads in BAM that are primary mapped** | 155,697,926<br>(98.64%) | 246,855,226<br>(98.91%) | 158,311,418<br>(98.57%) |
| **No. (proportion) of reads in BAM that are primary mapped and in proper pair** | 150,602,998<br>(95.41%) | 219,365,154<br>(87.90%) | 137,844,400<br>(85.83%) |

#### Variant calling and filtration for individuals

The number of invariant sites and SNPs called in the stock gDNA and amplified dilution datasets, and the proportion of sites retained in each dataset after filtration, are presented in Figure 3. The proportion of invariant sites (Fig 3A) and SNPs (Fig 3B) retained after filtration varied linearly across datasets, with the greatest proportion of sites retained in the unamplified stock gDNA dataset and the lowest proportion retained in the lowest-concentration dilution. Across all datasets, quality filters removed a greater proportion of heterozygous SNPs as compared to homozygous SNPs (Fig 3B).

**Figure 3.**
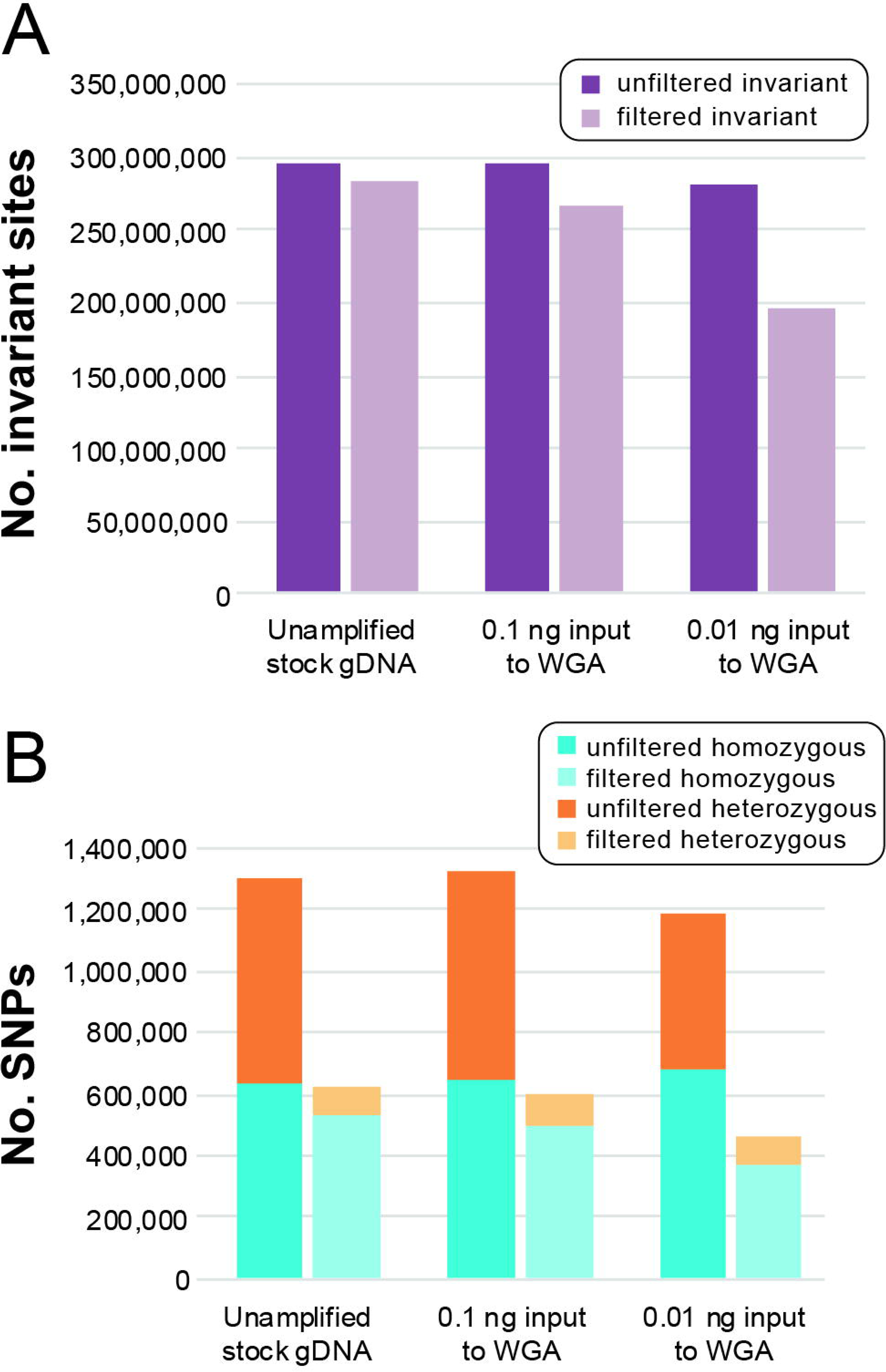
Number of sites in polymorphism datasets for the unamplified stock gDNA of an adult male of *Ancylostoma ceylanicum* and its whole genome amplified (WGA) serial dilutions. **(A)** Number of invariant sites before and after filtration. **(B)** Number of homozygous and heterozygous single nucleotide polymorphisms (SNPs) before and after filtration.

#### Genotype concordance and principal component analysis

Sensitivity, specificity, false discovery rate (FDR), and genotype concordance values for the amplified dilution datasets as compared against the unamplified stock gDNA “truth” dataset are summarized in Table 2. The number of sites available for comparison to the truth dataset varied by dilution, with the greater number of comparisons made in the higher-concentration dilution (290,351,177 sites total; 751,250 SNPs), and fewer comparisons made in the lower-concentration dilution (288,669,228 sites total; 763,384 SNPs; Table 2). The higher-concentration dilution had greater sensitivity (99.24%), specificity (>99.99%) and genotype concordance (97.07%), and a lower FDR (2.19%), as compared to the lower-concentration dilution, though specificity was similarly high (>99%) for both datasets (Table 2). Genetic similarity across all *A. ceylanicum* datasets, including outgroup L3 specimens, is visualized via PCA in Figure 4. Based on 747 SNPs retained after joint site-level and PLINK2 filters, principal components 1 and 2 explained 18.9% and 12.6% of variance in the dataset, respectively (31.5% cumulative variance). Points representing the unamplified stock gDNA truth dataset and the higher-concentration amplified dilution (0.1 ng to input to WGA) are plotted in a similar position relative to one another and the outgroup L3s, while the lower-concentration dilution (0.01 ng to input to WGA) is farther removed from the truth dataset along both principal component axes (Fig 4).

**Figure 4.**
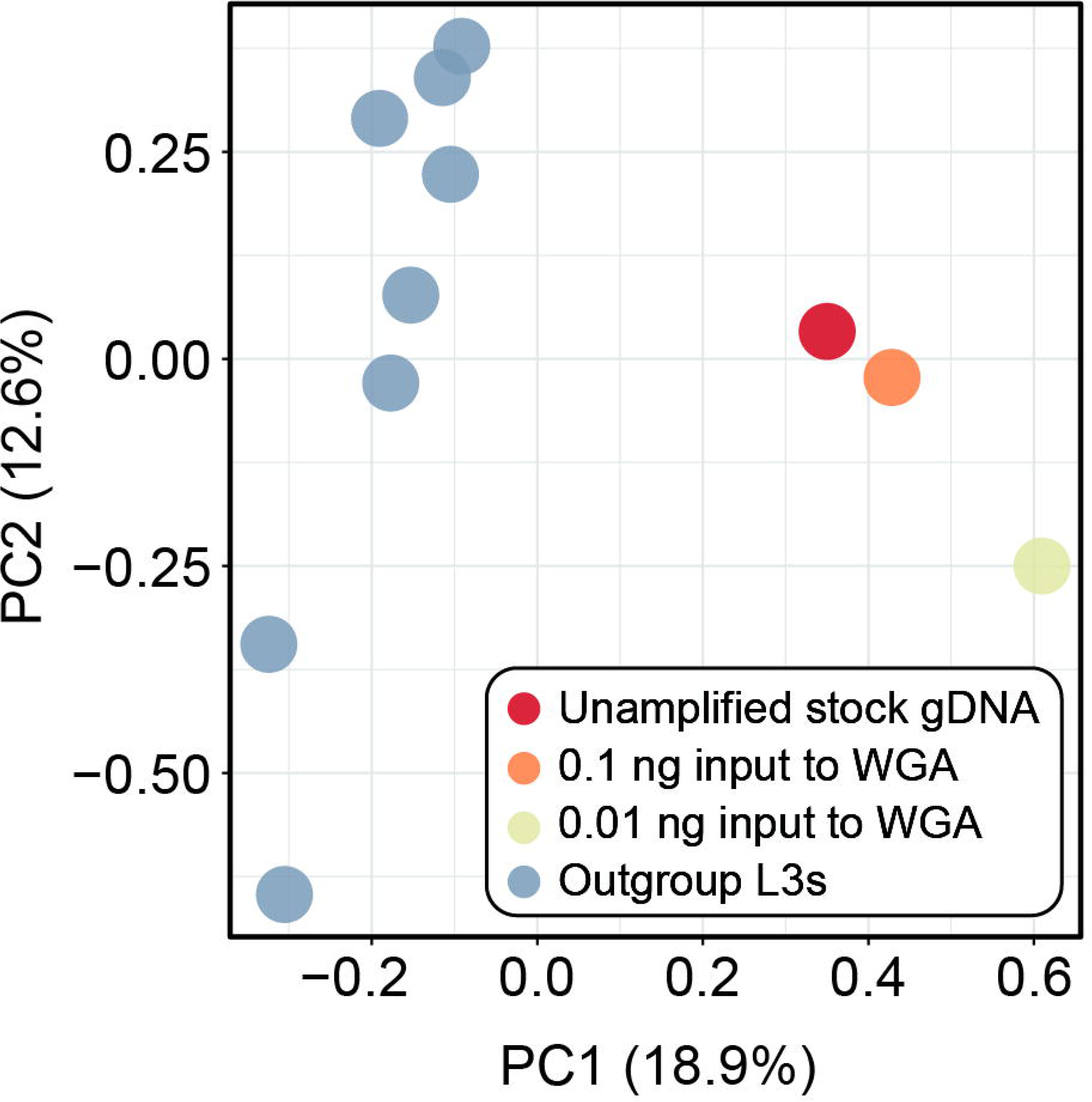
Principal component analyses (PCA) generated by PLINK2 from single nucleotide polymorphism (SNP) datasets for *Ancylostoma ceylanicum* based on 747 SNPs. Prior to analysis, datasets were jointly filtered for mininum minor allele frequency >0.05, <30% site missingness, linkage disequllibrium, and to retain only biallelic sites.

**Table 2.** Genotype concordance statistics for *Ancylostoma ceylanicum* for filtered genotype call datasets generated from serial dilutions of genomic DNA input to whole genome amplification as compared to the unamplified “truth” dataset generated using the same source genomic DNA. Sensitivity, specificity, false discovery rate, and genotype concordance are calculated based on sites where a call was made and retained post-filtration in both datasets.

|  | Input to WGA |  |
| --- | --- | --- |
|  | 0.1 ng | 0.01 ng |
| **Total no. sites compared against stock gDNA “truth” dataset** | 290,351,177 | 288,669,228 |
| **No. invariant sites** | 289,599,927 | 287,905,844 |
| **No. SNPs** | 751,250 | 763,384 |
| **Sensitivity** | 99.24% | 97.00% |
| **Specificity** | >99.99% | 99.99% |
| **False discovery rate** | 2.19% | 5.94% |
| **Genotype concordance** | 97.07% | 91.21% |
| **TN** | 257,364,775 | 187,058,802 |
| **TP** | 472,687 | 316,741 |
| **FN** | 43 | 113 |
| **FP** | 43 | 1,022 |
| **TP + TN** | 17,644 | 10,633 |
| **TP + FN** | 756 | 515 |
| **TP + FP** | 1,362 | 7,955 |
| **FP + FN** | 8 | 14 |
| **TN + FN** | 2,953 | 9,720 |
| **FP + TN** | 9,601 | 12,211 |
| **FP + TN + FN** | 14 | 14 |
| **Genotype call in either “truth” or “call” dataset only** | 32,481,291 | 101,251,488 |

### Application of validated approach–*Necator americanus*

#### Genomic DNA extraction from individual third-stage larvae

Cycle threshold values for entire volumes of extracted and bead-purified nucleic acid from individual *N. americanus* L3s are plotted against the standard curve in Figure 5. Using the extraction approach optimized in *A. ceylanicum*, a total mass of 0.02–0.13ng (0.07 ± 0.03; n=30) of gDNA per L3 was obtained. Following this confirmation of extraction efficiency, ITS qPCR screening then informed 23 additional *N. americanus* L3 extractions as being of sufficiently high concentration (C_t_<25.2) for input to WGA and library preparation. These include year 2019 field-collected L3s from Beposo, Ghana used to initiate a novel laboratory strain of *N. americanus* (F0s; n=3), year 2024 L3s from the fourteenth passage of the novel laboratory strain (F14s; n=13), and year 2024 L3s from additional human subjects in Beposo (BH10s; n=7) (see Table 3).

**Figure 5.**
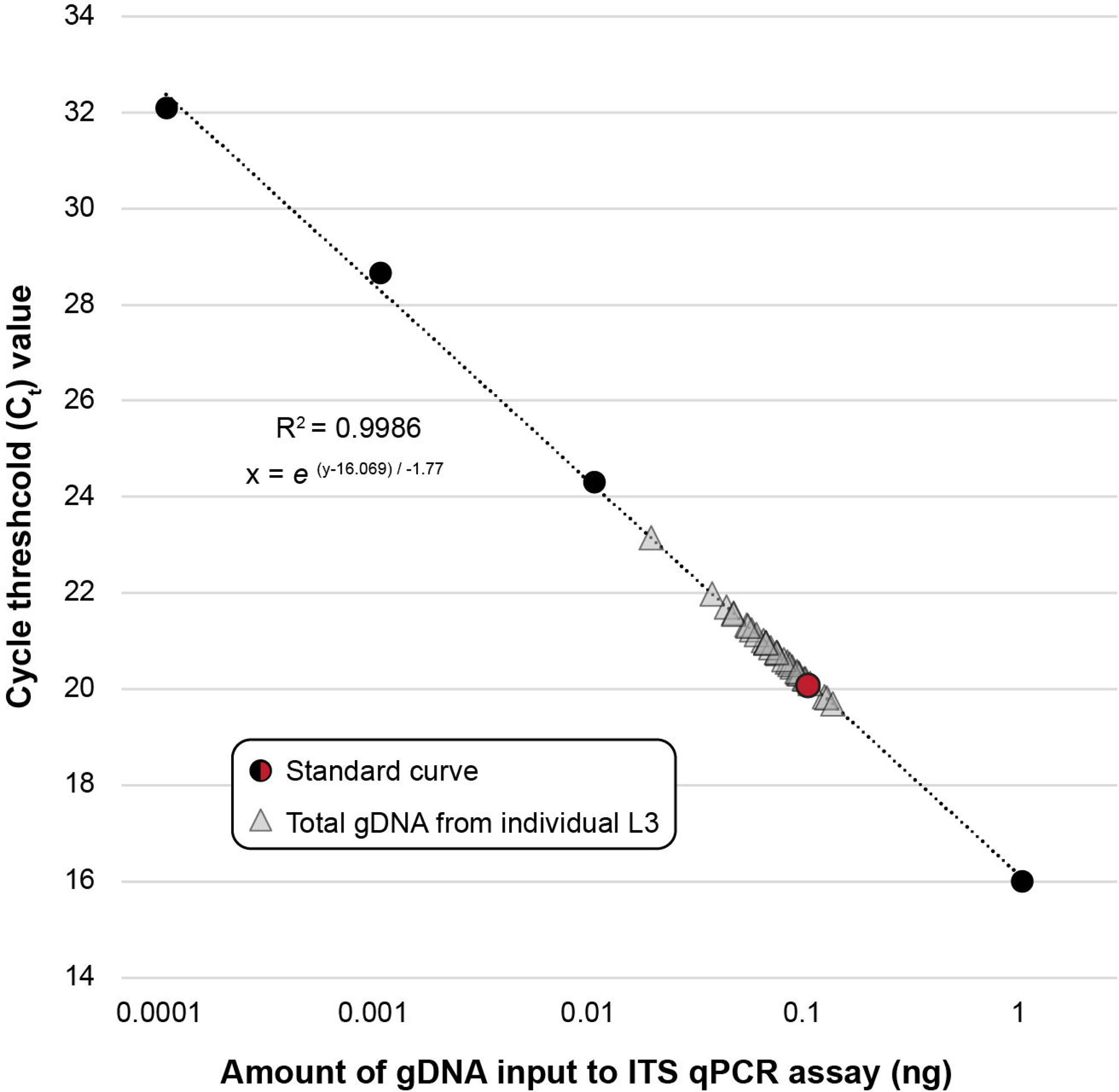
Cycle threshold (Ct) values for total volumes of genomic DNA (gDNA) extracted from individual third stage (L3) larvae of *Necator americanus.* Standard curve generated from serially diluted genomic DNA from a single adult male worm. Assay based on ITS2 primers of Llewellyn et al. (2016). Red point on standard curve indicates 0.1 ng.

**Table 3.** Whole genome amplification (WGA) and library preparation metrics for individual third-stage (L3) larvae of *Necator americanus* from the novel laboratory animal model (F14) and from human subjects in Beposo, Ghana (F0, BH10).

| Sample | Generation | Surface sterilized | Input to qPCR | Mean $C_t$ or $C_t$ | WGA time | Total mass post-WGA | Mass input to library prep | No. cycles indexing PCR |
| --- | --- | --- | --- | --- | --- | --- | --- | --- |
| Na L3 16 | F14 2024 | no | 1 μL in duplicate | 22.92 | 1 hr | 146.4 ng | 50 ng | 6 |
| Na L3 21 | F14 2024 | no | 1 μL in duplicate | 23.46 | 1 hr | 320 ng | 50 ng | 6 |
| Na L3 22 | F14 2024 | no | 1 μL in duplicate | 23.58 | 1 hr | 776 ng | 50 ng | 6 |
| Na L3 23 | F14 2024 | no | 1 μL in duplicate | 23.40 | 1 hr | 296 ng | 50 ng | 6 |
| Na L3 24 | F14 2024 | no | 1 μL in duplicate | 23.66 | 1 hr | 140.8 ng | 50 ng | 6 |
| Na L3 26 | F14 2024 | no | 1 μL in duplicate | 23.78 | 1 hr | 70.4 ng | 50 ng | 6 |
| Na L3 28 | F14 2024 | no | 1 μL in duplicate | 23.77 | 1 hr | 60.8 ng | 50 ng | 6 |
| Na L3 29 | F14 2024 | no | 1 μL in duplicate | 22.97 | 1 hr | 54.4 ng | 50 ng | 6 |
| Na L3 192 | F14 2024 | yes | 1 μL singular | 24.74 | 1.5 hr | >2,400 ng | 100 ng | 5 |
| Na L3 195 | F14 2024 | yes | 1 μL singular | 24.83 | 1.5 hr | 2,160 ng | 100 ng | 5 |
| Na L3 196 | F14 2024 | yes | 1 μL singular | 24.92 | 1.5 hr | 318 ng | 100 ng | 5 |
| Na L3 198 | F14 2024 | yes | 1 μL singular | 24.06 | 1.5 hr | >2,400 ng | 100 ng | 5 |
| Na L3 207 | F14 2024 | yes | 1 μL singular | 25.19 | 1.5 hr | 2,080 ng | 100 ng | 5 |
| Na L3 147 | F0 2019 | yes | 1 μL singular | 24.07 | 1.5 hr | 240 ng | 95 ng | 5 |
| Na L3 154 | F0 2019 | yes | 1 μL singular | 23.84 | 1.5 hr | 404 ng | 95 ng | 5 |
| Na L3 160 | F0 2019 | yes | 1 μL singular | 23.91 | 1.5 hr | 452 ng | 95 ng | 5 |
| Na L3 168 | BH10 2024 | yes | 1 μL singular | 23.71 | 1.5 hr | 720 ng | 95 ng | 5 |
| Na L3 169 | BH10 2024 | yes | 1 μL singular | 24.09 | 1.5 hr | 1096 ng | 95 ng | 5 |
| Na L3 178 | BH10 2024 | yes | 1 μL singular | 23.81 | 1.5 hr | >2,400 ng | 95 ng | 5 |
| Na L3 179 | BH10 2024 | yes | 1 μL singular | 23.97 | 1.5 hr | >2,400 ng | 95 ng | 5 |
| Na L3 181 | BH10 2024 | yes | 1 μL singular | 24.36 | 1.5 hr | 2,120 ng | 95 ng | 5 |
| Na L3 182 | BH10 2024 | yes | 1 μL singular | 24.19 | 1.5 hr | 158 ng | 95 ng | 5 |
| Na L3 190 | BH10 2024 | yes | 1 μL singular | 23.84 | 1.5 hr | 940 ng | 95 ng | 5 |

#### Next-generation sequencing, reference mapping, and variant filtration

The number of reads generated and mapped to the reference genome for each whole genome amplified *N. americanus* L3 are summarized in Table 4. While the total number of high-quality reads retained post-quality filtration varied somewhat across samples (210,448,984–504,066,846; mean 334,712,682), the proportion retained was relatively consistent (97.74–99.04%; mean 98.7%). The number and proportion of reads that mapped to the reference genome in primary alignment and in proper pair varied more considerably, however, with consistently higher mapping rates for F14s (56.5–90.88%; mean 77.23%; n=13) and BH10s (73.52–89.64%; mean 86.01%; n=7) than for F0s (33.64–64.16%; mean=45.13%; n=3). This in turn resulted in a greater proportion of the reference genome covered at >10× DOC for F14s (79.76–97.54%; mean 95.42%; n=13) and BH10s (63.06–97.71%; mean 89.11%; n=7) as compared to F0s (12.29–41.47%; mean=29.15%; n=3). These mapping and coverage rates correlated positively with the number of SNPs retained in individual datasets following site filtration, with a greater number of high-quality SNPs retained for F14s (2,444,496–3,127,840; mean 2,878,538; n=13) and BH10s (1,791,035–2,644,443; mean 2,470,267; n=7) as compared to F0s (357,926–1,301,134; mean 887,215; n=3) (see Table 4).

**Table 4.** Next-generation sequencing, reference mapping, and genotype calling metrics for individual third-stage (L3) larvae of *Necator americanus* from the novel laboratory animal model (F14) and human subjects in Beposo, Ghana (F0, BH10). Reads were aligned to a custom high-quality hybrid reference assembly generated from a single adult male worm from the first generation passage through the animal model.

|  | Sample | Surface sterilized | No. reads generated | No. reads retained post-QC | Percent retained | No. reads primary mapped and properly paired in BAM | Percent primary mapped and properly paired | Percent reference covered at >10x | No. SNPs post-filtration |
| --- | --- | --- | --- | --- | --- | --- | --- | --- | --- |
| F14 2024 | Na L3 16 | no | 387,338,376 | 382,056,588 | 98.64% | 242,596,740 | 63.50% | 96.34% | 2,858,909 |
|  | Na L3 21 | no | 504,066,846 | 497,350,222 | 98.67% | 339,314,806 | 68.22% | 97.21% | 2,866,177 |
|  | Na L3 22 | no | 343,076,466 | 338,890,744 | 98.78% | 268,364,630 | 79.19% | 97.31% | 2,822,592 |
|  | Na L3 23 | no | 428,421,268 | 422,658,010 | 98.65% | 301,436,350 | 71.32% | 97.54% | 2,668,364 |
|  | Na L3 24 | no | 415,493,452 | 410,809,630 | 98.87% | 280,564,390 | 68.30% | 96.75% | 2,870,844 |
|  | Na L3 26 | no | 410,478,580 | 405,192,074 | 98.71% | 330,245,484 | 81.50% | 96.28% | 3,075,356 |
|  | Na L3 28 | no | 446,454,728 | 440,868,248 | 98.75% | 249,083,724 | 56.50% | 96.28% | 2,919,114 |
|  | Na L3 29 | no | 348,664,668 | 343,595,388 | 98.55% | 254,234,324 | 73.99% | 96.82% | 2,928,072 |
|  | Na L3 192 | yes | 318,993,100 | 315,783,746 | 98.99% | 278,187,850 | 88.09% | 96.39% | 3,127,840 |
|  | Na L3 195 | yes | 340,301,672 | 337,035,200 | 99.04% | 303,287,396 | 89.99% | 96.87% | 2,993,041 |
|  | Na L3 196 | yes | 330,036,984 | 326,657,452 | 98.98% | 279,230,758 | 85.48% | 79.76% | 2,444,496 |
|  | Na L3 198 | yes | 299,724,208 | 296,692,880 | 98.99% | 269,647,064 | 90.88% | 97.35% | 2,918,875 |
|  | Na L3 207 | yes | 250,566,736 | 247,597,576 | 98.82% | 215,608,256 | 87.08% | 95.50% | 2,927,310 |
| F0 2019 | Na L3 147 | yes | 279,455,188 | 276,154,932 | 98.82% | 92,901,956 | 33.64% | 12.29% | 357,926 |
|  | Na L3 154 | yes | 322,168,330 | 317,417,496 | 98.53% | 119,334,176 | 37.60% | 33.70% | 1,002,585 |
|  | Na L3 160 | yes | 338,364,048 | 330,727,252 | 97.74% | 212,201,306 | 64.16% | 41.47% | 1,301,134 |
| BH10 2024 | Na L3 168 | yes | 302,810,796 | 299,190,264 | 98.80% | 268,188,116 | 89.64% | 91.16% | 2,569,602 |
|  | Na L3 169 | yes | 213,124,946 | 210,448,984 | 98.74% | 184,341,866 | 87.59% | 89.15% | 2,585,924 |
|  | Na L3 178 | yes | 329,549,428 | 324,536,320 | 98.48% | 287,404,646 | 88.56% | 97.46% | 2,565,561 |
|  | Na L3 179 | yes | 275,442,906 | 270,930,184 | 98.36% | 237,193,060 | 87.55% | 97.71% | 2,644,443 |
|  | Na L3 181 | yes | 252,352,266 | 249,380,952 | 98.82% | 221,248,550 | 88.72% | 96.41% | 2,628,279 |
|  | Na L3 182 | yes | 339,498,884 | 335,383,758 | 98.79% | 246,565,652 | 73.52% | 63.06% | 1,791,035 |
|  | Na L3 190 | yes | 222,007,800 | 219,060,190 | 98.67% | 189,396,496 | 86.46% | 88.81% | 2,507,027 |

#### Principal components analysis and maximum likelihood inference

Genetic similarity across *N. americanus* SNP datasets is visualized via PCA in Figure 6. For the dataset containing only the F14 and BH10 specimens, which comprised 4,635 SNPs after joint site-level and PLINK2 filters, principal components 1 and 2 explained 24.4% and 10.4% of the variance in the dataset, respectively (34.8% cumulative variance)––similar to the proportion of variance explained in the *A. ceylanicum* validation dataset. In this analysis, specimens clearly group by population of origin, with F14s representing a tighter, more cohesive cluster than BH10s, which are more broadly distributed across the second axis of variation (see Fig. 6A). For the analysis including the additional three F0 specimens, which comprised 3,276 SNPs after joint site-level and PLINK2 filters, principal components 1 and 2 explained 25.5% and 11.4% of variance in the dataset, respectively (36.9% cumulative variance). In this analysis, F14s again form a clearly defined cluster, with BH10s and F0s demonstrating more variability along the second axis of variation. BH10s and F0s did not overlap with one another along the first axis of variation, and their separation from one another was less distinct than from F14s (see Fig. 6B). Maximum likelihood (ML) topologies as inferred by RAxML are presented in Figure 7. For the dataset containing only F14 and BH10 specimens (911 sites analyzed), the results of ML inference reflect those of PCA, with the two populations recovered as separate clades with strong support, and longer branch lengths inferred for BH10 specimens as compared to the F14s (see Fig. 7A). For the dataset including F0s (379 sites), F14s were again recovered as a clade, with BH10s and F0s forming a separate clade with relatively longer branches (see Fig. 7B). With this clade of field-collected samples, two subclades were recovered: one containing four of seven BH10 specimens, and one containing the remaining BH10 specimens intermixed with F0 specimens (see Fig. 7B). In both topologies, clades separating lab and field-collected samples were well-supported by ultrafast bootstrap (BS) values, with relatively low support (BS <70) for most internal nodes between individuals within a clade (see Fig. 7).

**Figure 6.**
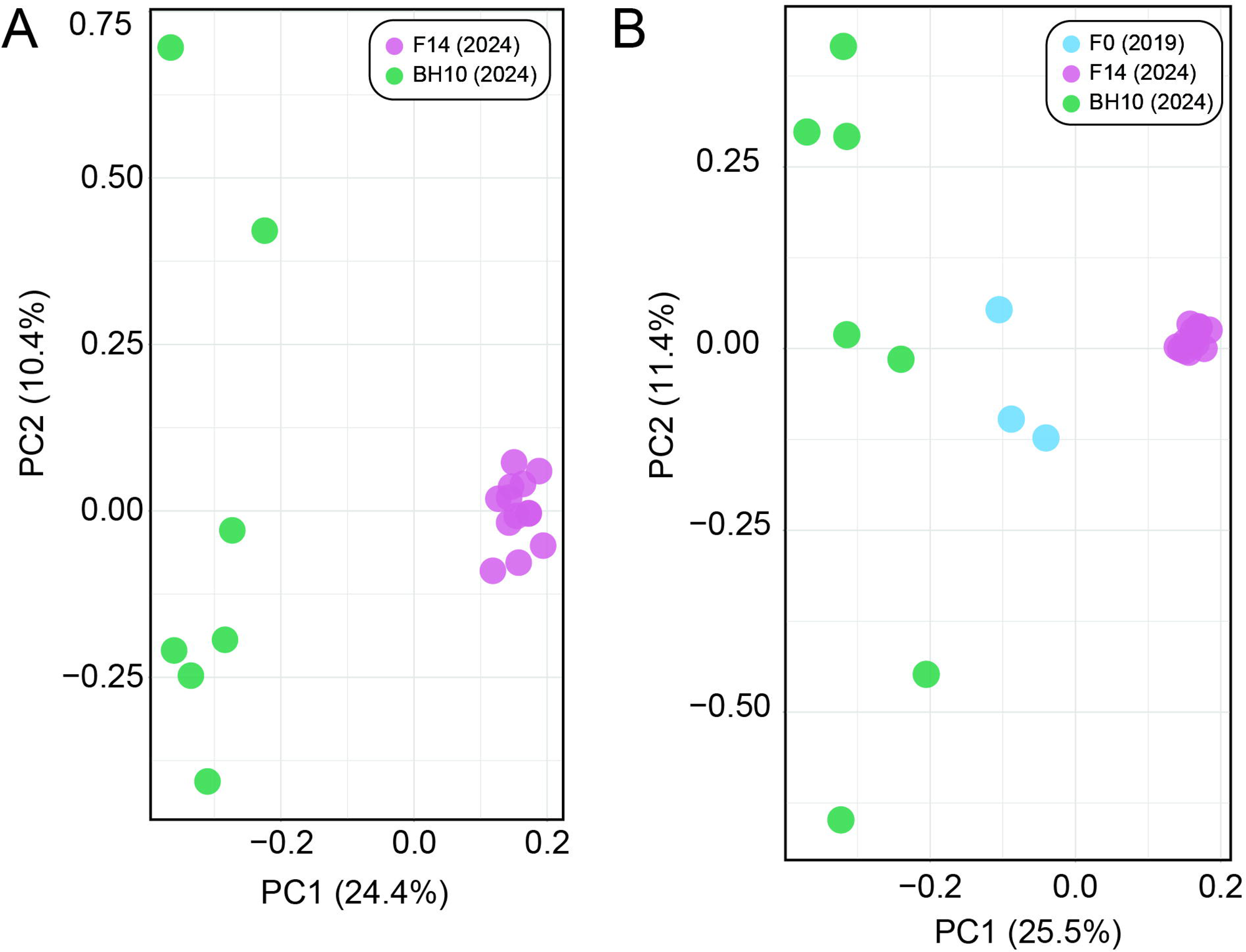
Principal component analyses (PCAs) generated by PLINK2 from single nucleotide polymorphism (SNP) datasets for Necator americanus. A. Dataset with only F14s and BHlOs based on 4,635 SNPs. B. Full dataset including FOs based on 3,276 SNPs. Prior to analysis, both datasets were jointly filtered for mininum minor allele frequency >0.05, <30% site missingness, linkage disequllibrium, and to retain only biallelic sites.

**Figure 7.**
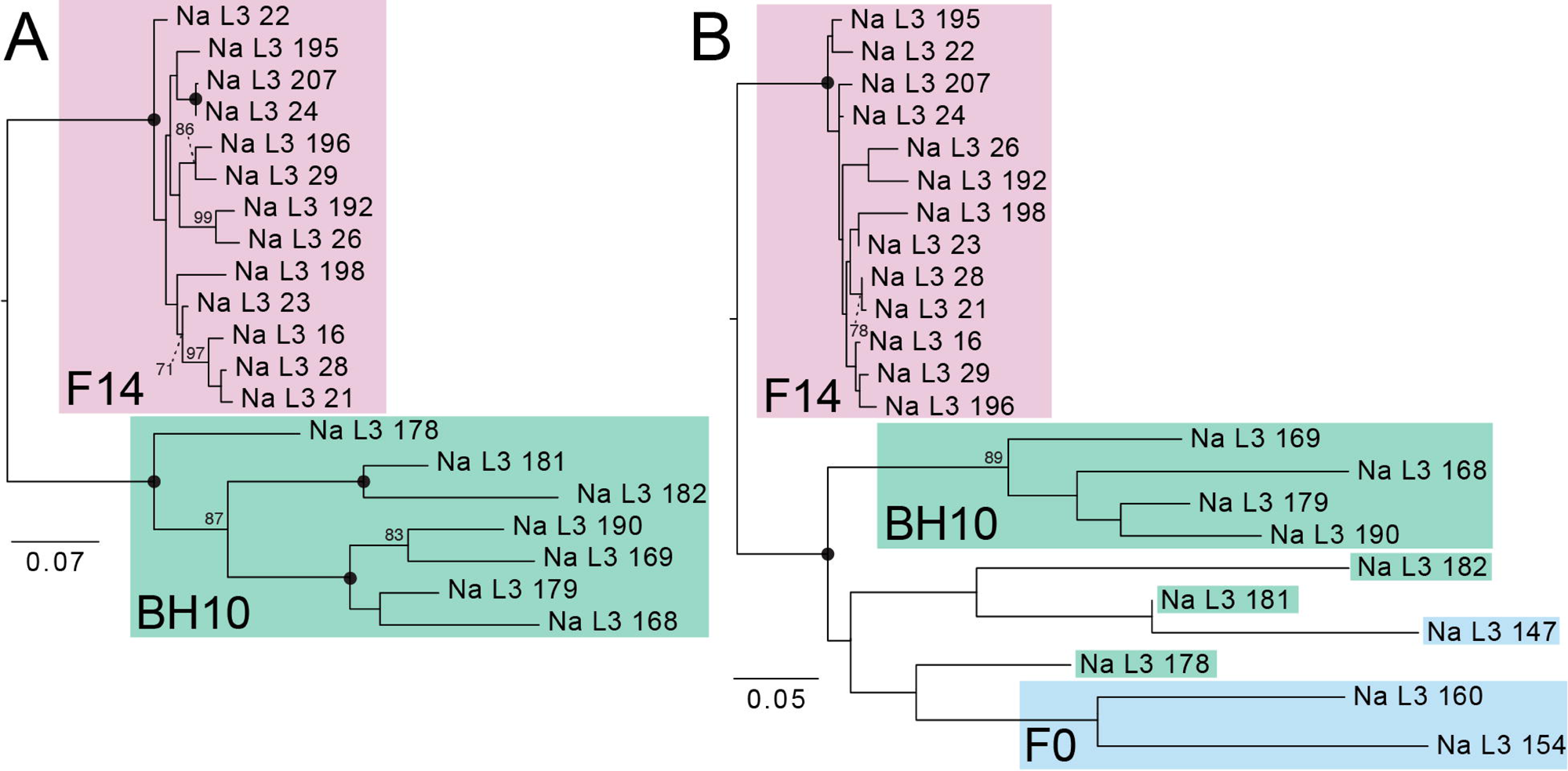
Maximum liklihood trees generated by RAxML from single nucleotide polymorphism (SNP) datasets for *Necator americanus* rooted to maximize subtree balance. Prior to analysis, both datasets were jointly filtered to retain sites with a minor allele frequency >0.05 and <30% site missingness and to remove invariant sites. **A.** Dataset with only F14s and BHl0s based on 911 sites. B. Full dataset including F0s based on 379 sites. Nodal support values are generated from 1,000 rapid bootstrap (BS) replicates; solid circles represent BS values of 100; BS values with low support <70 are not shown. Scale bars at left indicate nucleotide substitutions per site.

#### Proportion of heterozygous sites in laboratory versus field-collected samples

The number and proportion of heterozygous SNPs in individual filtered VCFs for F14s and BH10s are summarized in Table 5. The proportion of heterozygous sites ranged from 9.90–22.07% (15.29% ± 3.48%; n=13) for F14s and 13.65–26.37% (21.33% ± 3.85%; n=7) for BH10s. A one-tailed student’s t-test indicates that BH10s have on average a significantly greater proportion of heterozygous SNPs as compared to F14s (p-value=0.001). It is worth noting that the BH10 specimen with the fewest number of total SNPs retained post-filtration (Na L3 182; 1,791,035 SNPs) also had the lowest proportion of heterozygous SNPs of any BH10 specimen (13.65%; see Table 5). BH10s also had a greater estimated nucleotide diversity as compared to F14s (π of 0.27585 vs. 0.1093) and a greater number of private alleles (40,803 vs. 29,006; see Table 5).

**Table 5.** Number and proportion of heterozygous sites in *Necator americanus* SNP datasets for specimens from the 14^th^ generation of the novel laboratory animal model (F14) from 2024 and human subjects in Beposo, Ghana (BH10) from 2024. A one-tailed student’s t-test assuming homoskedasticity (p=0.001) supports that BH10s have a statistically greater proportion of heterozygous SNPs as compared to F14s.

| | Sample | Total no. SNPs | No. het. SNPs | Proportion of SNPs that are het. | Average proportion of het. SNPs | $\pi$ | No. private alleles |
| --- | --- | --- | --- | --- | --- | --- | --- |
| F14 2024 | Na L3 16 | 2,858,909 | 551,380 | 19.29% | 15.29%<br>($\pm 3.48\%$) | 0.1093 | 29,006 |
|  | Na L3 21 | 2,866,177 | 400,281 | 13.97% |  |  |  |
|  | Na L3 22 | 2,822,592 | 453,857 | 16.08% |  |  |  |
|  | Na L3 23 | 2,668,364 | 588,899 | 22.07% |  |  |  |
|  | Na L3 24 | 2,870,844 | 503,867 | 17.55% |  |  |  |
|  | Na L3 26 | 3,075,356 | 340,317 | 11.07% |  |  |  |
|  | Na L3 28 | 2,919,114 | 510,652 | 17.49% |  |  |  |
|  | Na L3 29 | 2,928,072 | 464,618 | 15.87% |  |  |  |
|  | Na L3 192 | 3,127,840 | 309,580 | 9.90% |  |  |  |
|  | Na L3 195 | 2,993,041 | 368,184 | 12.30% |  |  |  |
|  | Na L3 196 | 2,444,496 | 285,629 | 11.68% |  |  |  |
|  | Na L3 198 | 2,918,875 | 437,323 | 14.98% |  |  |  |
|  | Na L3 207 | 2,927,310 | 482,560 | 16.48% |  |  |  |
| BH10 2024 | Na L3 168 | 2,569,602 | 537,906 | 20.93% | 21.33%<br>($\pm 3.85\%$) | 0.27585 | 40,803 |
|  | Na L3 169 | 2,585,924 | 548,946 | 21.23% |  |  |  |
|  | Na L3 178 | 2,565,561 | 575,624 | 22.44% |  |  |  |
|  | Na L3 179 | 2,644,443 | 610,740 | 23.10% |  |  |  |
|  | Na L3 181 | 2,628,279 | 693,181 | 26.37% |  |  |  |
|  | Na L3 182 | 1,791,035 | 244,531 | 13.65% |  |  |  |
|  | Na L3 190 | 2,507,027 | 541,403 | 21.60% |  |  |  |

## DISCUSSION

### Development of a novel workflow for consistent nucleic acid extraction from individual third-stage hookworm larvae

Due to their size and accessibility from both laboratory animals and human subjects, L3s are arguably the most feasible stage in the hookworm life-cycle to target for individual whole genome sequencing. Both aims of this study (validation and application) thus depended entirely on the ability to extract quantifiable amounts of nucleic acid from individual L3s. Therefore, we first developed an optimized approach that delivers consistent and usable concentrations of nucleic acid in the range of 0.02–0.13ng total (0.07 ± 0.03; see Fig. 5). To our knowledge, this is the first published workflow optimized specifically for purifying nucleic acid from single hookworm L3s. Here, we leveraged this approach for WGA, NGS, and SNP discovery. However, nucleic acid purification is use case agnostic, and therefore this same approach can be applied to generate input for a variety of amplification-based approaches in which an individual hookworm, rather than a pooled sample, would be desired. This includes, for example, qPCR assays aimed at identifying a locus linked to a phenotype of interest, or targeted amplicon-based sequencing approaches like that of George et al. [22] to investigate drug tolerance. We further demonstrate the efficacy of qPCR screening using species-specific ITS primers to measure the concentration of individual extractions using small inputs (1µL of nucleic acid; see Fig. 1, Table 3). As variation in extraction efficiency was observed across specimens (see Fig. 5, Table 3) this screening approach proved crucial for identifying which extractions of *N. americanus* L3s could be reliably used as input to WGA.

### Surface sterilization and sample preservation impact sequence quality and reference mapping rates in field-collected samples

The way specimens were fixed and treated prior to nucleic acid extraction impacted the quality and quantity of read-level alignments and resulting SNP datasets in largely predictable ways. For example, F14s and BH10s were preserved in RNAlater, which prevents nucleic acid degradation over long-term storage. These specimens collectively shared high rates (mean=80.3%) of reads mapped to the reference genome in primary alignment and in proper pair, covering most of the reference genome (mean=93.21%) at >10× DOC, and producing SNP datasets with numerous (mean=~2.7 million) high-quality genotype calls post-filtration (see Table 4). By contrast, F0s, which were older (collected in 2019 versus 2024), preserved in 10mM Tris, and underwent several freeze-thaw cycles prior to extraction, had appreciably lower mapping (mean=45.13%) and coverage (mean=29.15%) rates, and fewer SNPs (mean=~887,000) retained in final datasets as compared to their better-preserved counterparts. Given that the reference genome for *N. americanus* used here was generated from a single adult male from the first passage through the animal model, it is unlikely that low mapping rates in F0s are the result of divergence from the reference, especially considering F14s and BH10s demonstrate high mapping rates. Given that similar numbers of reads were generated across samples, this indicates that the difference lies in the proportion of reads generated from each sample that are hookworm gDNA. The GenomiPhi V3 Ready-To-Go DNA Amplification Kit used here relies on multiple displacement amplification (MDA) to replicate the gDNA in a sample, and this process has been demonstrated to be biased against fragmented DNA, as increased fragmentation results in fewer priming sites on template strands, and thus fewer opportunities for hyperbranching [23]. Therefore, any amount of non-hookworm gDNA in a sample, even if small in concentration, has the potential to preferentially amplify during WGA if it is less fragmented than the hookworm gDNA. This could be expected with, for example, bacterial genomes, which have a comparably small, circularized structure that is highly stable [24]. In fact, investigatory metagenomic analysis using the CZ ID platform [25] indicated a high proportion of bacterial and viral reads in F0 datasets. This discrepancy motivated us to generate two final SNP datasets for *N. americanus*–one with, and one without, the F0 specimens. More broadly, our results suggest that for poorly preserved L3s input to WGA, surface sterilization cannot alone overcome the bias of MDA against fragmented DNA, highlighting the importance of proper nucleic acid preservation.

### Input to whole genome amplification impacts breadth and depth of genome coverage for low-input sample types

The goal of the validation aim of this study was to determine whether genome-wide SNP datasets can be reliably generated from low-concentration gDNA from individual L3s. To do so, we leveraged easily-accessible specimens of *A. ceylanicum* from a long-maintained laboratory strain [26,27]. Informed by the results of our nucleic acid extraction test for L3s of both *A. ceylanicum* and *N. americanus* (see Figs. 1 & 5, respectively), we serially diluted nucleic acid from a single adult male (1ng/µL) tenfold (to 0.1ng/µL and 0.01ng/µL) to represent the range of concentrations observed for individual L3s. A 1µL aliquot of each dilution was subjected to WGA and then sequenced alongside the unamplified stock gDNA, which serves as a comparative truth dataset. Our results clearly indicate that for low input sample types (the manufacturer’s recommended lower limit for input to a GenomiPhi V3 reaction is 10ng), WGA introduces bias in both the breadth and depth of coverage distributions across the genome, but that these metrics are more similar to unamplified samples when higher concentrations of gDNA are input to amplification (see Fig. 2). In terms of breadth, as input to WGA decreases, additional reads are needed to reach the same depth of coverage obtained in the unamplified truth dataset. For the lower-concentration dilution (0.01ng input to WGA), coverage plots suggest a plateau around 70–75%, indicating it may not even be possible to achieve equivalent breadth of coverage in very low input hookworm sample types using WGA (see Fig. 2A). Our results suggest this breadth of coverage bias is likely due to partiality in which regions of the genome are replicated during WGA. This is evidenced by depth of coverage plots for each dataset, in which amplified samples show significant proportions of the genome at coverage levels far exceeding those achieved in the unamplified truth dataset (see Fig. 2B). When considering only sites at or below 70× DOC, the higher-concentration dilution dataset closely resembles the coverage profile of the unamplified truth dataset, while the lower-concentration dilution covers comparatively fewer sites to depths <70×. The MDA technology on which the GenomiPhi V3 kit is based is known to preferentially amplify some genomic regions based on, for example, GC content or conformational characteristics, and preferential amplification of some regions over others due simply to random chance (i.e., drift-based bias) is also an inherent risk [23]. Unevenness in coverage distributions has been reported previously when applying WGA in other systems, including for the model nematode, *Caenorhabditis elegans* [28]. Ultimately, characterization of this bias in our system informed our decision to forgo use of a maximum depth filter in SNP filtration, as for WGA samples, this would remove high-quality sites that are covered at great depth due not to alignment error, but rather the nature of the approach. In fact, implementing a maximum depth cutoff of <100× reduced values of sensitivity, specificity, and genotype concordance for both amplified dilutions. In all, we demonstrate that WGA introduces bias in breadth and depth of coverage when sequencing individual L3s, and that bias can be mitigated by increasing both sequencing effort and the amount of gDNA input to WGA; however, increases in sequence effort have diminishing returns at very low-concentration inputs. It is worth noting that *A. ceylanicum* validation datasets were aligned to the reference genome of Schwarz et al. [29] which was generated using worms from a different laboratory strain than those sequenced here. Inbreeding and drift in these isolated systems, which are a well-documented consequence of laboratory models [30], or misassembles in this reference, which was generated using only short-read data, therefore have the potential to explain the plateau at ~94% breadth of coverage for the unamplified dataset.

### Optimized filtration scheme removes a greater proportion of heterozygous SNPs as compared to homozygous SNPs

Our results demonstrate that, across *A. ceylanicum* validation datasets, a greater proportion of heterozygous SNPs were removed during site-level filtration as compared to homozygous SNPs or invariant sites (see Fig. 3). When considering SNPs only, filtration for minimum quality score, minimum read depth, and biallelism collectively removed similar proportions of homozygous and heterozygous sites across datasets (~10%, 15%, and 37% of each type of SNP removed in the unamplified truth, 0.1ng input, and 0.01ng datasets, respectively; see Fig. 3B). Enforcement of a strand bias <0.5 filter then removed ~20%, ~33%, and ~44% of remaining homozygous SNPs in the unamplified truth, 0.1ng input and 0.01ng input datasets, respectively, but removed ~75% of remaining heterozygous SNPs across all datasets (Fig. 3B). Strand bias is defined as an error of short-read sequencing platforms where one strand in a read pair is preferred over the other, which can misrepresent support for the alleles present, resulting in an incorrect genotype call [31]. The goal of strand bias filtration is therefore to reduce false positive SNP calls [32]. In developing an optimal filtration scheme for WGA from L3s, we noted that filtering on minimum quality score, minimum read depth, and biallelism alone produced suboptimal metrics for genotype concordance and FDR (~87% and ~8%, respectively) in the highest-concentration dilution dataset. Further investigation revealed that the majority of miscalls involved an error at only one of two alleles at a site (i.e., only very few miscalls were the result of a homozygous reference site [0/0] being called a heterozygous alternate site [1/1] or vice versa). Characterizing this error profile led us to implement a strict strand bias filter. Though its implementation did result in the removal of some correctly-called sites (mostly homozygous alternate [TP] and heterozygous [TP/TN] SNPs) the concomitant reduction in miscalled sites greatly improved concordance metrics. Allelic dropout, or the loss of an allele due to its homolog being preferentially amplified, is a known potential consequence of MDA, particularly with very low inputs [33]. It is not possible to determine from this experimental design whether the patterns we observed at heterozygous sites in WGA datasets result from strand bias during sequencing or artifacts of MDA, or a combination thereof. Despite not knowing its exact origins, our results demonstrate that a strict strand bias filter effectively mitigates the impact of this error profile in downstream comparative analyses.

### Amplification bias results in high levels of missingness across samples

Following standard practice, individual VCFs filtered at the site level were merged and jointly filtered for minimum minor allele frequency and a cutoff for site-level missingness prior to PCA and ML tree building. This joint filtration drastically reduced the number of SNPs in all three datasets (the *A. ceylanicum* validation dataset, and the two *N. americanus* datasets with and without F0s). For the *A. ceylanicum* validation dataset, joint filtration removed ~98% of sites. For the *N. americanus* datasets, joint filtration removed ~98.8% of sites in the F14 + BH10 dataset, and ~99.6% of sites for dataset including F0s. Further investigation into the *A. ceylanicum* validation dataset revealed that amplified dilutions and amplified outgroup L3s had 60–70% missing data across sites in the joint VCF (see Supplemental Figure 1). It is important to note that all specimens in the *A. ceylanicum* SNP dataset had similar SNP counts after individual filtration (431,583–684,083; mean=581,556 SNPs). Therefore, high proportions of missingness across samples after joint filtration suggest that while similar number of high-quality SNPs are retained for each individual, where these high-quality calls occur in the genome often does not overlap between samples. Based on our previous observations of coverage discrepancies introduced by WGA, we suggest this is likely the result of drift-based amplification bias, and perhaps to a lesser extent, our strict filtration criteria. In other words, there is likely some randomness in which genomic regions get amplified during WGA and retained post-filtration for each sample. Despite this, we still retained a sufficient number of jointly-called sites across samples to achieve resolution in PCA and ML trees. This highlights a need to develop a targeted sequencing approach for hookworm population genomics, akin to the genome-wide amplicon-based MAD^4^HatTeR assay of Aranda-Días et al. [34] for *Plasmodium*. Designing a similar approach in hookworms will require intensive research and development, as the sites of interest to be included in a hookworm genome-wide amplicon-based sequencing panel (e.g., for identifying population structure, drug tolerance, etc.) have been only minimally investigated. These could include, for example, the β-tubulin gene, which is implicated in drug tolerance [35,36] and *COX1*, which has been previously used for hookworm population genetics [10,11]. In the absence of an amplicon panel, other researchers conducting population genomics from small input sample types have shown combining WGA with restriction-site associated DNA sequencing (RAD-seq) as a promising alternative approach [37,38]. After WGA, digestion with a restriction enzyme and selection of only fragments containing those enzyme binding sites for library preparation should ensure that a greater proportion of sequencing effort per sample is dedicated to sites that will also be present in other samples, mitigating drift-based amplification bias. Until such time as the development of an amplicon-based sequencing assay for hookworm population genomics becomes tractable, this is a future avenue worth exploring to improve the resolution of the methodology described here.

### Input to whole genome amplification impacts variant call accuracy and interpretations of relatedness

Our genotype concordance results for *A. ceylanicum* validation datasets demonstrate that input to WGA is positively correlated with not only evenness of breadth and depth of genome coverage, but also variant call number and accuracy. While sensitivity was only marginally higher in the higher-concentration dilution as compared to the lower-concentration dilution (~99% versus 97% respectively), genotype concordance and FDR differed appreciably between the two datasets (~97% and ~2%, respectively, in the higher-concentration dilution versus ~91% and ~5%, respectively, in the lower-concentration dilution; see Table 2). The similarly high rates of specificity in both datasets are attributable to the large number of TN sites (homozygous reference [0/0]) called across all datasets. Genotype concordance data also demonstrate the challenge of missingness in WGA datasets and its negative correlation with input, with >32 million sites (0.1ng input to WGA) and >101 million sites (0.01ng input to WGA) incomparable due to a genotype call missing in either the “truth” or the dilution dataset (see Table 2). Nevertheless, a high genotype concordance value and a low FDR for the higher-concentration dilution indicates that, despite the biases introduced by WGA, highly accurate variant call datasets can be generated for amplified samples given sufficient input of nucleic acid to WGA and strict SNP filtration. Furthermore, our results from quantifying entire volumes of nucleic acid extracted from individual *N. americanus* L3s demonstrate that, although variation in concentration exists across samples, we can reliably purify ≥0.1 ng from individual L3s (see Fig. 5). This ultimately informed our C_t_ value cutoff for qPCR screening in *N. americanus* extractions, ensuring only samples with a total mass of ~0.1ng or greater were input to WGA, library preparation, and sequencing. A high genotype concordance and a low FDR in the higher-concentration dilution predictably translated to a nearly-identical plotting of this dilution and the unamplified stock gDNA “truth” dataset on the PCA, which was not the case for the lower-concentration dilution (see Fig. 4). This indicates that, even with reduction in the number of SNPs available for comparison as a result of joint filtration and further thinning for linkage disequilibrium, accurate estimates of relatedness and population structure can still be obtained using amplified samples. We are not the first to validate the use of WGA for helminth population genomics. The significant work of Choi et al. [21] in identifying recrudescence versus reinfection in filarial worms informed our validation methodology. However, there is a key difference between our two approaches. Choi et al. used gDNA from multiple adult males of *Brugia malayi* as input to WGA to inform use of their approach with individual microfilariae, but they did not dilute the nucleic acid from these adults prior to WGA and sequencing. We believe we have here taken a more rigorous approach to validation, whereby gDNA from an adult male hookworm was diluted down to the concentrations observed for extractions from individual L3s prior to WGA and sequencing. We then compared these amplified dilution datasets against the same unamplified stock gDNA from which they were generated, ensuring our ability to fully describe the biases associated with WGA when using inputs well below the manufacturer’s recommended >10ng input.

### Population assignment and genomic similarity for laboratory versus field-collected samples

Applying our validated approach, we compared year 2024 L3s from the fourteenth passage through a novel laboratory strain of *N. americanus* (F14s) to year 2024 L3s from human subjects in Beposo, Ghana (BH10s). Importantly, this is the same hookworm endemic region from which L3s collected in 2019 were used to establish the novel laboratory strain (F0s; Harrison, Herzog et al. 2025). Results from PCA and ML inference for F14s and BH10s clearly support the two groups as distinct populations, with more genomic diversity encompassed by BH10s as inferred by greater distance between points in PCA (Fig. 6A), longer branch lengths in ML topologies (see Fig. 7A), a greater number of private alleles, and a greater estimated value of π (Table 5). Results for datasets including a small number of F0 specimens were similar between the two approaches, with F14s recovered as a more genetically homogenous population separate from field-collected samples (see Figs. 6B, 7B). Unfortunately, high proportions of missingness in F0s (likely due to poor preservation, see above) limited their utility in comparative analyses. That high proportions of missingness can result in incorrect interpretations of relatedness in PCA and ML inference are well-documented phenomena [39,40]. Therefore, we cannot have the same confidence in our assessment of genomic similarity and population assignment for these samples. Nevertheless, results from PCA and tree building, which indicate F0s as more similar to other field-collected samples from the same endemic community (BH10s) than to later passages through the animal model (F14s), align with expectations (see Figs. 6B, 7B). This further highlights the importance of sample preservation considerations when using bias-prone methodologies such as WGA.

### Heterozygosity in laboratory versus field-collected samples

We found that in filtered individual-level SNP datasets, F14s had on average significantly fewer heterozygous SNPs as compared to their BH10 field-collected counterparts (see Table 5). The results from our validation aim show that while strand bias filtration does remove more heterozygous SNPs than it does homozygous SNPs, the proportion of heterozygous SNPs removed is consistent across datasets (see above, and Fig. 3). Therefore, we are confident that comparing the proportion of total SNP sites that are heterozygous in a sample after filtration still allows for unbiased comparison of overall proportions of heterozygous sites between the two populations. These results demonstrate that reduction in heterozygosity is detectable in the laboratory strain five years after initiation of infection, but that individual worms still retain numerous heterozygous sites despite multiple generations of serial passage. To our knowledge, this is the first explicit quantification of heterozygosity in a laboratory strain of human hookworms. Reduced heterozygosity is a signature of inbreeding and genetic drift, which are expected to occur through time in laboratory strains [41–44], and in many cases, this is even the desired outcome [45]. Given that most knowledge of the genetic dynamics of eukaryotic laboratory models comes from rodents, species of *Drosophila*, or *C. elegans*, these findings highlight a knowledge gap in our understanding of how non-model invertebrates, and particularly helminths, respond to laboratory rearing, and adds to the growing body of literature on the subject [46–48]. Though not an explicit goal of this experiment, it is valuable to note that the reference genome for *N. americanus* used here, which was assembled using nucleic acid from a single adult male hookworm from the first-generation passage through laboratory animals proved an equally suitable reference from which to call SNPs for both laboratory and field-collected samples. This is evidenced by similar mapping rates and numbers of SNPs called for both F14s and BH10s (see Table 4). Given the known importance of reference selection in short-read mapping [49] as well as the potential for genomic change caused by adaptation to laboratory animals and conditions [50], confirmation that the F1 reference genome is suitable for SNP calling in field-derived samples underscores the feasibility of continued exploration of population structure of *N. americanus* in endemic communities.

## MATERIALS AND METHODS

### Human subjects collections and animal care

Prior to enrolling human subjects, research study protocols were independently reviewed and approved by the IRB committees of Yale University and the Noguchi Memorial Institute for Medical Research at University of Ghana. Informed consent was obtained from all human study subjects enrolled in the field survey conducted in Beposo, Ghana. Animal husbandry was provided by the Yale Animal Resource Center and animal experimentation protocols approved by the Yale Animal Care and Use Committee prior to study initiation.

The human strain of *N. americanus* described here was originally isolated in 2019 from individuals (n=300) living in Beposo, which is located in the Pru West District in the Bono-East region of Ghana. Fecal samples from study subjects determined to be hookworm positive using Kato-Katz microscopy were mixed with bone charcoal (Ebonex, Melvindale, Michigan) and cultured at 25° C for 10–14 days. Viable third stage larvae (L3) of hookworm were collected and concentrated using a Baermann funnel apparatus. Species-specific PCR of cultured larvae confirmed monocultures of *N. americanus* [51,52].

*In vivo* laboratory propagation of the Beposo *N. americanus* strain was achieved through serial infection of weanling (21 days old) male Golden Syrian Hamsters of the outbred HsdHan: AURA (LVG) strain (Envigo, Indianapolis, IN). Prior to infections with *N. americanus* L3, hamsters (n=6 per cage group) were provided drinking water *ad libitum* containing dexamethasone (2.5µg/ml) for 4 days. Animals were then injected subcutaneously with 300 L3 suspended in PBS and monitored for fecal egg excretion using McMaster slide microscopy [53]. L3 obtained from hamster fecal cultures were used to initiate subsequent passages in weanling animals.

The BH10 isolates of human *N. americanus* L3 were cultured from subjects participating in an ongoing longitudinal study (2022–2025) of hookworm infection in Beposo. Fecal samples from infected study subjects (n=134) were pooled and cultured in a temporary field laboratory to obtain L3. Viable larvae were combined and resuspended in RNAlater prior to transport to the United States (Yale University and University of Nebraska Medical Center) for further study.

## Data availability

Quality-controlled sequence data generated as part of this study are deposited in the National Center for Biotechnology Information (NCBI) Sequence Read Archive (SRA) under the BioProject accession ID PRJNAXXXXXX and under BioSample accession nos. SAMNXXXXXX (*Ancylostoma ceylanicum*) and SAMXXXXXXX (*Necator americanus*).

### Approach validation–Ancylostoma ceylanicum

#### Dilution series and standard curve generation

Genomic DNA was extracted following the methods described by Herzog et al. [16] from a single adult male of *Ancylostoma ceylanicum* obtained from the laboratory strain maintained at the Yale School of Public Health and preserved in RNAlater (Thermo Fisher Scientific). The extraction was quantified via Qubit™ double-stranded high sensitivity DNA assay (dsDNA HS; Invitrogen) then serially diluted tenfold from 1ng/µL to 0.0001ng/µL using nuclease-free water. To form a standard curve against which extractions of individual L3s could be measured, quantitative PCR (qPCR) was used to generate cycle threshold (C_t_) values for each dilution. Samples were amplified in duplicate on a Mic qPCR Cycler (Bio Molecular Systems) using Luna® Universal qPCR Master Mix (New England Biolabs) and *A. ceylanicum*-specific forward and reverse primers for ITS1 [54] following the manufacturer’s recommended cycling protocol.

#### Optimization of genomic DNA extraction from individual third-stage larvae

L3s of *A. ceylanicum* were obtained from the laboratory strain maintained at the Yale School of Public Health and preserved in RNAlater (Thermo Fisher Scientific). Individual L3s were isolated in phosphate buffered saline (PBS; 1X; pH 7.4; Thermo Fisher Scientific) on a glass spot plate using an Olympus SZ40 dissecting microscope and a micropipette, added to 25µL of lysis buffer, and incubated at 55°C for 24hrs. Three types of lysis buffer were tested to determine which would produce the highest gDNA yield: the modified CGP lysis buffer of Choi et al. [21], the MagMAX™ Viral/Pathogen Nucleic Acid Isolation Kit lysis buffer (Thermo Fisher Scientific), and the Zymo Quick-DNA™ High Molecular Weight MagBead Kit lysis buffer (Zymo Research). Each lysis buffer was prepared according to the authors’ or manufacturers’ instructions, and eight L3s were extracted per buffer. Following lysis, extractions were purified using KAPA Pure Beads (Roche) in a 1:1 bead-to-sample ratio and eluted in 25µL nuclease-free water. All extractions were quantified via qPCR as described above and compared against the standard curve of serially diluted gDNA from the adult male.

#### Whole genome amplification (WGA)

To best-represent the range of gDNA concentrations observed from individual L3 extractions, the two highest-concentration dilutions of stock gDNA from the single adult male worm (0.1ng/µL and 0.01ng/µL) were chosen for input to WGA. The eight highest-concentration extractions of individual L3s were also amplified to serve as outgroup specimens in downstream comparative analyses. Samples underwent WGA using a GenomiPhi V3 Ready-To-Go DNA Amplification Kit (Cytiva). Either 1µL (for the two dilutions) or 2µL (for the eight L3s) of template gDNA was used as input to WGA and amplified for 1.5hr following the manufacturer’s recommendations. Amplified samples were then purified using KAPA Pure Beads (Roche) in a 2:1 bead-to-sample ratio, eluted in 20µL nuclease-free water, and quantified via Qubit dsDNA HS assay.

#### Library preparation and next-generation sequencing

The unamplified stock gDNA from the single adult male (1ng/µL), its amplified serial dilutions (0.1ng/µL and 0.01ng/µL), and the eight amplified L3 outgroup specimens were sent to the Yale Center for Genome Analysis (YCGA) for library preparation and next-generation sequencing. Libraries were prepared using an xGen™ cfDNA & FFPE DNA Library Preparation Kit (Integrated DNA Technologies) with 5–13 cycles of indexing PCR (see Table 1) and sequenced on an Illumina NovaSeq 6000 for 2×150 paired-end sequencing targeting 50× DOC. Reads were demultiplexed by the YCGA.

#### Variant calling

Demultiplexed reads were filtered for quality, length, and adaptor contamination using fastp v.0.23.2 [55] and read quality post-filtration was further assessed using FastQC v0.12.1 [56]. Quality-controlled reads were aligned to the *A. ceylanicum* reference genome [29] using BWA-MEM v.0.7.17 [57] with default settings. Resulting SAM files were converted to BAM files using samtools v.1.19.2 [58]. Optical duplicates were removed with the “MarkDuplicates” tool of the Genome Analysis Toolkit (GATK) v. 4.4.0.0 [31,59] specifying an optical pixel distance of 2500. For the unamplified stock gDNA dataset and those from its amplified serial dilutions, subsampled BAM files at 10-million-read increments were also generated using samtools [58]. Summary statistics for BAM files were generated using the “flagstats” and “stats” functions in samtools [58] and DOC was further summarized with the aid of MultiQC v.1.8 [60]. The “mpileup” command in bcftools v. 1.9 [61] was used to generate BCF files from BAM files, specifying the output of read depth, allele depth, and strand bias terms using the “--annotate FORMAT” flag. Variant call format (VCF) files were generated from BCF files file using the bcftools [61] “call” command, specifying use of the multiallelic caller to retain SNPs and discard insertions and deletions (indels) and invariant sites. For the unamplified stock gDNA dataset and those from its amplified serial dilutions, “call” was run a second time to generate VCFs containing both SNPs and invariant sites to be used for comparative genotype concordance analyses. The bcftools [61] “view” command was used to filter all resulting VCFs to retain only biallelic SNPs with a minimum quality score of 30, a minimum read depth of 10, and a maximum strand bias value of 0.5. The proportion of heterozygous SNPs in datasets before and after filtration was measured using bcftools [61].

#### Genotype concordance calculations and principal component analysis

The “GenotypeConcordance” tool in picard v.3.0.0 [59] was used to calculate genotype concordance metrics for the two amplified dilution datasets based on filtered VCFs containing both SNPs and invariant sites. The comparative “truth” dataset was specified as the filtered VCF for the unamplified stock gDNA containing SNPs and invariant sites (see Table 2). Custom bash scripts were used to calculate sensitivity, specificity, and FDR from the outputs of “GenotypeConcordance”. Filtered VCFs containing only SNPs generated from each dataset (including outgroup L3s) were then used as input to PCA in PLINK2 v2.00a1LM [62]. Individual VCFs were first combined into a joint VCF using bcftools [61]. The joint VCF was then filtered using vcftools v.0.1.16 [63] for minimum minor allele frequency of 0.05 and a maximum site-level missingness of 30%. Filters for linkage disequilibrium (using a window size of 50kb, a window step size of 10bp, and an r^2^ threshold of 0.1) and biallelism across samples were applied in PLINK2, and PCAs were visualized using ggplot2 v.3.4.1 [64] in R v.4.2.3 [65] via RStudio v.2023.06.1+524 [66].

### Application of validated approach–*Necator americanus*

#### Standard curve generation

Genomic DNA was extracted following the methods described by Herzog et al. [16] from a single adult male of *Necator americanus* preserved in RNAlater (Thermo Fisher Scientific) from the first passage (F1) through the animal model established and maintained at the Yale School of Public Health. The extraction was quantified and serially diluted tenfold from 1ng/µL to 0.0001ng/µL as described above for *A. ceylanicum*. To form a standard curve, qPCR was used to generate C_t_ values for each dilution. Samples were amplified in duplicate on a QuantStudio7 Pro thermal cycler (ThermoFisher Scientific) using Luna® Universal qPCR Master Mix (New England Biolabs) and *N. americanus*-specific forward and reverse primers for ITS2 [54] following the manufacturer’s recommended cycling protocol.

#### Genomic DNA extractions from individual third-stage larvae

Fourteenth generation L3s (year 2024; hereafter F14s) were obtained from the Ghana laboratory strain for *N. americanus*, preserved in RNAlater (Thermo Fisher Scientific), and stored at −80°C prior to processing. L3s from human subjects in Beposo, Ghana were also isolated for comparison. These included two populations: The first generation used to establish infection in the animal model (year 2019; hereafter F0s) and specimens collected in summer 2024 as part of a longitudinal epidemiologic study in Beposo (hereafter BH10s). These L3s were field collected as described above and preserved in 10mM Tris (F0s) or RNAlater (Thermo Fisher Scientific; BH10s) before being transported to the United States for storage at −80°C. Individual L3s from each of these three populations were manually isolated for gDNA extraction as described above for *A. ceylanicum*, individually added to 25µL of Zymo Quick-DNA™ High Molecular Weight MagBead Kit lysis buffer (Zymo Research) and incubated at 55°C for 24hrs. For a subset of L3s (see Table 3), surface sterilization was performed prior to lysis as follows: L3s were isolated in the well of a glass spot plate in 1X PBS, then PBS was replaced with hydrochloric acid (HCl; 1% v/v), and L3s were incubated in HCl for 5min at RT. After surface sterilization, HCl was removed and L3s were washed 3–5 times with 1X PBS. Following 24hr of lysis, extractions were purified using KAPA Pure Beads (Roche) in a 1:1 bead-to-sample ratio and eluted in 11µL nuclease-free water. Extractions were then quantified via qPCR as described above for dilutions of adult *N. americanus* gDNA and compared against the standard curve. For a subset of F14s, qPCR screening was run in duplicate, while for the remaining samples the assay was run singularly (see Table 3). For a separate set of a F14s (n=30), gDNA was eluted in 10µL following bead purification and the entire elution volume was input to a single qPCR reaction to measure the total mass of gDNA obtained from individual *N. americanus* L3s.

#### Whole genome amplification

Extractions with a Ct ≤25.2 were used as input to WGA (see Table 3). The GenomiPhi V3 Ready-To-Go DNA Amplification Kit (Cytiva) was used with the entire volume remaining from each extraction after qPCR screening (9–10µL) as input, and 1–1.5hr of amplification (see Table 3). Amplified samples were then purified using KAPA Pure Beads (Roche) in a 2:1 bead-to-sample ratio, eluted in 20µL nuclease-free water, and quantified via Qubit dsDNA HS assay.

#### Library preparation, next-generation sequencing and variant calling

Short-read sequencing libraries were generated from amplified gDNA of *N. americanus* L3s in-house using a DNA Prep Tagmentation Kit (Illumina) following the manufacturer’s protocol with 50–100ng of amplified gDNA as input and 5–6 cycles of indexing PCR (see Table 3). Libraries were quantified via Qubit dsDNA HS assay and average insert sizes were measured via TapeStation D1000 High Sensitivity DNA ScreenTape analysis (Agilent Technologies). Libraries were then sent to the YCGA where they were pooled by concentration and further size-selected via gel electrophoresis and excision prior to 2×150 paired-end sequencing on an Illumina NovaSeq 6000 targeting 70× depth of coverage per sample. Reads were demultiplexed by the YCGA. Read-level data were then quality controlled and variant call datasets were generated following the approach outlined above for *A. ceylanicum* and using a custom hybrid assembly from a single adult male collected from the animal model as a reference genome. Unlike for *A. ceylanicum* however, VCFs containing invariant sites were not generated for *N. americanus*, as genotype concordance analyses were not necessary for this dataset.

#### Principal component analysis and maximum likelihood phylogenetic inference

Filtered VCFs for individual *N. americanus* L3s were combined into joint VCFs and filtered jointly as described above for *A. ceylanicum*. Two joint VCFs were generated: one with all amplified *N. americanus* L3s (n=23) and one with only the F14 and BH10 specimens (n=20). Joint VCFs were used as input to PCA and ML inference. For PCA, joint filtration and analysis followed the approach outlined above for *A. ceylanicum*. For ML inference, joint VCFs filtered for minimum minor allele frequency of 0.05 and a maximum site missingness of 30% were converted to PHYLIP files and invariant sites were removed using the Python 3 script *raxml_ascbias* (ascbias.py; https://github.com/btmartin721/raxml_ascbias#raxml_ascbias) in Python v.3.9.7 [67]. Methods for ML inference using RAxML v.8.2.12 [68] follow Herzog et al. [69]. The “populations” module in STACKS v2.68 [70,71] was used to calculate nucleotide diversity (π) and the number of private alleles from the jointly filtered dataset containing only F14s and BH10s.

## ACKNOWLEDGEMENTS

We are grateful to the residents of Beposo and surrounding communities in Ghana for their generous participation in this study. We also thank the field, laboratory and administrative teams at the Noguchi Memorial Institute for Medical Research and the University of Ghana for their collaboration. We thank Dr. Richard Clopton and Deb Clopton (Peru State College) for aid in development of techniques for individual L3 isolation, and Dr. John Hawdon (George Washington University) for conveying the technique used for surface sterilization. We also thank Dr. Michael Wiley and Dr. Mara Jana Broadhurst (University of Nebraska Medical Center) for use of select laboratory equipment. This work was supported by National Institutes of Health National Institute of Allergy and Infectious Diseases award numbers R01AI182301, R01AI162826 and UL1TR001863, and by a Nebraska Research Initiative award. Research reported in this publication was supported by the National Institute of General Medical Sciences of the National Institutes of Health under award number 1S10OD030363-01A1. This work was completed utilizing the Holland Computing Center of the University of Nebraska, which receives support from the UNL Office of Research and Innovation, and the Nebraska Research Initiative.

