## Supplementary figures and images for "Hookworm genomic diversity and population structure from accessible sample types: A validated approach to generate genome-wide polymorphism datasets from individual third-stage larvae"

### Supplemental Figure 1

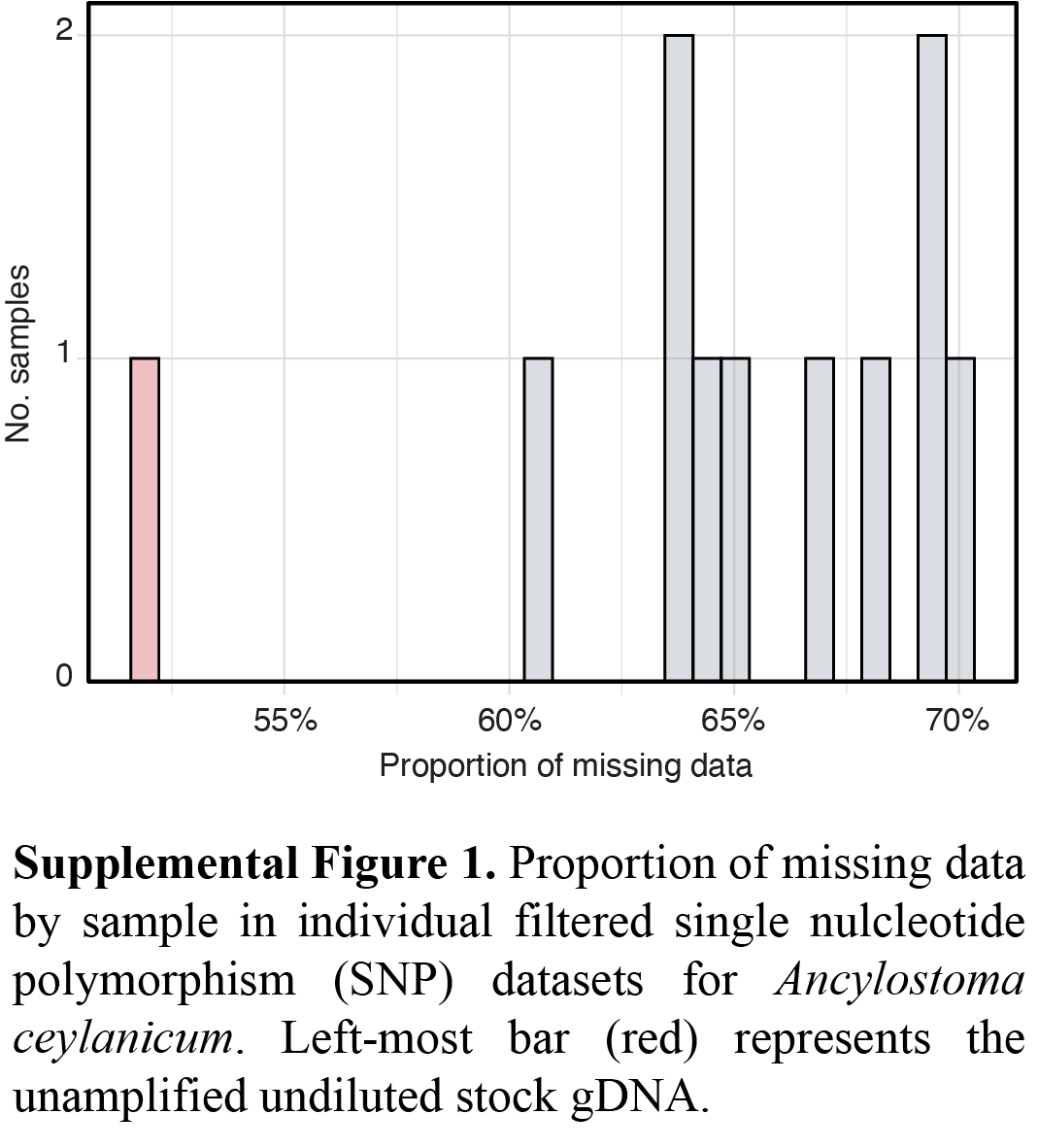
